# Conditional approach as cooperation in predator inspection: a role for serotonin?

**DOI:** 10.1101/436345

**Authors:** Ana Flávia Nogueira Pimentel, Tamires dos Santos Carvalho, Fernando Lima, Monica Lima-Maximino, Marta Candeias Soares, Caio Maximino

**Affiliations:** Laboratório de Neurociências e Comportamento “Frederico Guilherme Graeff”, Universidade Federal do Sul e Sudeste do Pará, Marabá/PA, Brazil; Faculdade de Psicologia, Universidade Federal do Sul e Sudeste do Pará, Marabá/PA, Brazil; Laboratório de Neurofarmacologia e Biofísica, Universidade do Estado do Pará, Marabá/PA, Brazil; CIBIO, Centro de Investigação em Biodiversidade e Recursos Genéticos, Universidade do Porto, Vairão, Portugal

**Keywords:** Serotonin, Cooperation, Conditional approach, Predator inspection, Guppy

## Abstract

In guppies (*Poecilia reticulata*), a small number of individuals break away from a shoal and approach a potential predator, a behavior termed “predator inspection”. These animals often employ a “conditional approach” strategy, in which an individual approaches the predator in the first move and subsequently approaches it only if a second individual swims even with it during inspection. This strategy is analogous to the “tit-for-tat” strategy of the Prisoner’s Dilemma, suggesting that it could be used to study cooperation. Serotonin is thought to mediate cooperative behavior in other fish species. Exposure to the animated image of a predator in a tank that contained a parallel mirror – mimicking an equally cooperating conspecific – promoted inspection and decreased refuge use, but increased freezing, suggesting that conditional approach is also associated with fear. To understand whether serotonin participates in conditional approach in guppies, we treated animals with either vehicle (Cortland’s salt solution), fluoxetine (2.5 mg/kg) or metergoline (1 mg/kg), and tested then in a predator inspection paradigm. Fluoxetine increased the time the animal spent inspecting the predator image, while metergoline decreased it. Fluoxetine also decreased time spent avoiding the predator and increased freezing, while metergoline decreased freezing. These results suggest that phasic increases in serotonin levels promote conditional approach, suggesting a role for this neurotransmitter in cooperation.

Preprint: https://doi.org/10.1101/436345; Data and scripts: https://github.com/lanec-unifesspa/TFT

## 1. Introduction

Sometimes being seen is better than blending into the crowd. And when it comes to interactions between putative preys and their predators, it may not be enough to stay vigilant as to simply detect a predator more rapidly, but one also needs to signal its presence and awareness (e.g. head bobbing and fin flicking [1]) and sometimes even to provide information regarding own physical condition (e.g. leaping in artiodactyls [2]). Indeed, predators seem to be less successful when prey are fully aware of their presence, in part due to the improved vigilance effect in which groups are able to detect predators sooner than solitary individuals [3]. One seemingly paradoxical behaviour occurs when some of these prey leave the shoal to slowly approach and inspect the predator presumably to reduce the uncertainty of the situation and to be able to gain valuable information to all other conspecifics involved [4]. Since this behaviour implies costs to the individual(s) that interacts with the predator but contributes to the fitness of the shoal, it has been interpreted as cooperative, however, not without strategic specificities.

Conditional approach is a behavioural strategy used by different fish species, including guppies *Poecilia reticulata* [5–8] and sticklebacks *Gasterosteus aculeatus* [9,10], and acts as an incentive for predator inspection, given that the risk of approaching a predator is shared between all inspectors, but not with the individuals that keep a distance [11,12]. Conditional approach represents a cooperative strategy that is comparable to a Prisoner’s Dilemma game [13,14], in which cheating a partner (by staying behind) is the most profitable option while joint inspection is the most beneficial (symmetrical) form of cooperation, allowing the advantages of inspecting the predator to overcome those of remaining at a safe distance [11]. Trivers [15] proposed the Prisoner’s Dilemma as a mechanism to interpret the evolution of reciprocal altruism, and Axelrod and Hamilton [14] showed that “tit-for-tat” is an evolutionary stable strategy. This strategy instructs the player to cooperate in the first move and to copy subsequent moves by the partner. Tit-for-tat requires a pre-specified payoff matrix that is, in the case of conditional approach, unknown [5]; however, conditional approach is analogous to tit-for-tat in that when fish inspect in pairs, due to the higher risk of taking the lead, exchanges in leading position are an example of cooperation based on reciprocity [5,6]. Prisoner’s Dilemma therefore models a situation in which each individual receives a worse pay-off for not cooperating than for cooperating [12,14,15]. In conditional approach, non-cooperating individuals also obtain information regarding the predator without incurring in risk, but mutual reciprocal cooperation offers animals a better pay-off. Conditional approach is only employed by inspecting individuals in cooperative partnerships; predator inspection *per se* can be made by solitary individuals, as has been shown for guppies [16,17] and zebrafish (*Danio rerio*) [18,19]. Predator inspection by singletons is still cooperative, as the rest of the shoal can benefit from the transmission of information about the predator, but involves inspection only by one individual [17,20].

The guppy is a small ovoviviparous fish originating from Central America and northern South America [21]. It is well known by its bright colours, and widely used in ornamental aquaria. Seghers [22] observed that some guppies, when facing a predator at a distance in the wild, soon inspect this potential threat. Guppies leave the shoal individually or in small groups, and approach the potential threat to obtain information. There is evidence that guppies use conditional approach during this inspection behaviour [5–7,20,23,24]. Dugatkin [5] suggested that conditional approach presents the basic characteristics of tit-for-tat: it is “nice” (that is, the first “move” is to cooperate by initiating inspection), “retaliatory” (animals defect by retreating from inspection if the conspecific does not reciprocate), and “forgiving” (animals cooperate by re-initiating inspection if a previously defecting conspecific changes its course and begins inspection).

Serotonin (5-hydroxytryptamine, 5-HT) is a monoamine that is synthesized from tryptophan. This monoamine participates in the modulation of stress and defensive behaviour in fish [25–27], and has been implicated in vertebrate social behaviour [27,28]. Converging evidence from human and non-human animal research suggest that altering 5-HT levels can directly influence social perception and mood, with decreased serotonin leading to isolation and decreased sociality [29]. Moreover, it has been shown that 5-HT participates directly in prosocial and cooperative behaviour: tryptophan depletion reduces cooperation in the Prisoner’s Dilemma in human participants, and decreases the probability of cooperative responses given previous mutually cooperative behaviour [30]. In the cleaner wrasse *Labroides dimidiatus,* acute treatment with fluoxetine or the 5-HT_1A_ receptor agonist (±)-8-Hydroxy-2-1A (dipropylamino)tetralin (8-OH-DPAT) increased client inspection and tactile stimulation, while treatment with the tryptophan hydroxylase inhibitor para-chlorophenylalanine or the 5-HT 1A receptor antagonist N-[2-[4-(2-Methoxyphenyl)-1-piperazinyl]ethyl]-N-2-pyridinylcyclohexanecarboxamide (WAY 100,635) decreased client inspection and cleaners’ cheating levels [31]. WAY 100,635 also delays learning of client value, suggesting that this receptor is involved in the perception of danger and therefore in how cleaners appraise, acquire information and respond to client-derived stimuli [32].

Given that predator inspection entails the evaluation of the level of predator threat, as well as of the value of cooperating vs. defecting with conspecifics, it is probable that 5-HT is involved in the mediation of conditional approach/cooperative behaviour in guppies. We modified the method described by Dugatkin [5], using a video animation of a sympatric predator (the blue acará cichlid *Aequidens pulcher)* presented to a single guppy in a tank with a parallel mirror. We predicted that the predator animation, compounded with the parallel mirror – mimicking an equally cooperating conspecific, with approaches and retreats to the predator –, would elicit predator approach (inspection), leading animals to spend more time in the inspection section than near the predator rather than in the avoidance zone or in the refuge. We also predicted that the predator animation would elicit fear-like behaviour, including freezing, as well as higher time spent in the avoidance zone and in the refuge. The first prediction was confirmed, but the second prediction was only partially true, given that the animated image increased freezing, but not time in the avoidance zone, and decrased time in the refuge zone. We also tested the hypothesis that serotonin promotes conditional approach by treating animals with fluoxetine or metergoline, a non-selective serotonin receptor antagonist. We predicted that phasic 5-HT (increased by acute fluoxetine) would promote conditional approach, while simultaneously increasing fear/anxiety (as observed in zebrafish: [33]); blocking receptors with metergoline (thereby decreasing the role of tonic 5-HT) would decrease conditional approach and decrease fear. The first prediction was confirmed, but metergoline only decreased freezing, without affecting conditional approach.

## 2. Methods

### 2.1. Animals, housing, and baseline characteristics

For the experiments of the present research, 46 male guppies were used. Animals were bought from a commercial vendor, and left to acclimatize to the laboratory (LaNeC) for two weeks before beginning experiments. The animals were bred at the vendor, and represent the third generation from populations captured in the wild in the Amazon (Belém/PA, Brazil). These domesticated animals represent a mix of strains. Animals were fed daily with commercial feed. The animals were housed collectively in 40 L tanks, at a maximum of 25 fish per tank, with the following water quality parameters being controlled: dissolved oxygen ≈7.8 mg / L, ammonia <0.002 mg/L, pH 7.0 – 8.0, temperature 24-30 °C.

### 2.2. Induction of conditional approach

A glass tank (40.6 cm x 18 cm x 25 cm) was used, containing an artificial plant in one extreme (“refuge zone”) and a computer screen (Samsung T20c310lb, 20”, LED screen, nominal brightness 20 cd/m^3^) positioned on the opposite extreme. A mirror was positioned in parallel to one of the sides of the tank for the whole duration of experiments. The avoidance zone is defined as the half of the tank that is farthest from the screen, minus the refuge zone, and the inspection zone is defined as the other half of the tank (Figure 1). The avoidance zone, therefore, did not include the refuge. While the mirrored image does not respond in the way a live individual would (i.e. there is no exchange of the leading position, which is the position entailing more risk), and therefore the mirrored image can only be as cooperative as the focal individual, using a mirror to simulate cooperation has been shown to work and in fact reflect the cooperativeness of a specific individual, despite the limitations in mimicking a natural behavior [5,9].

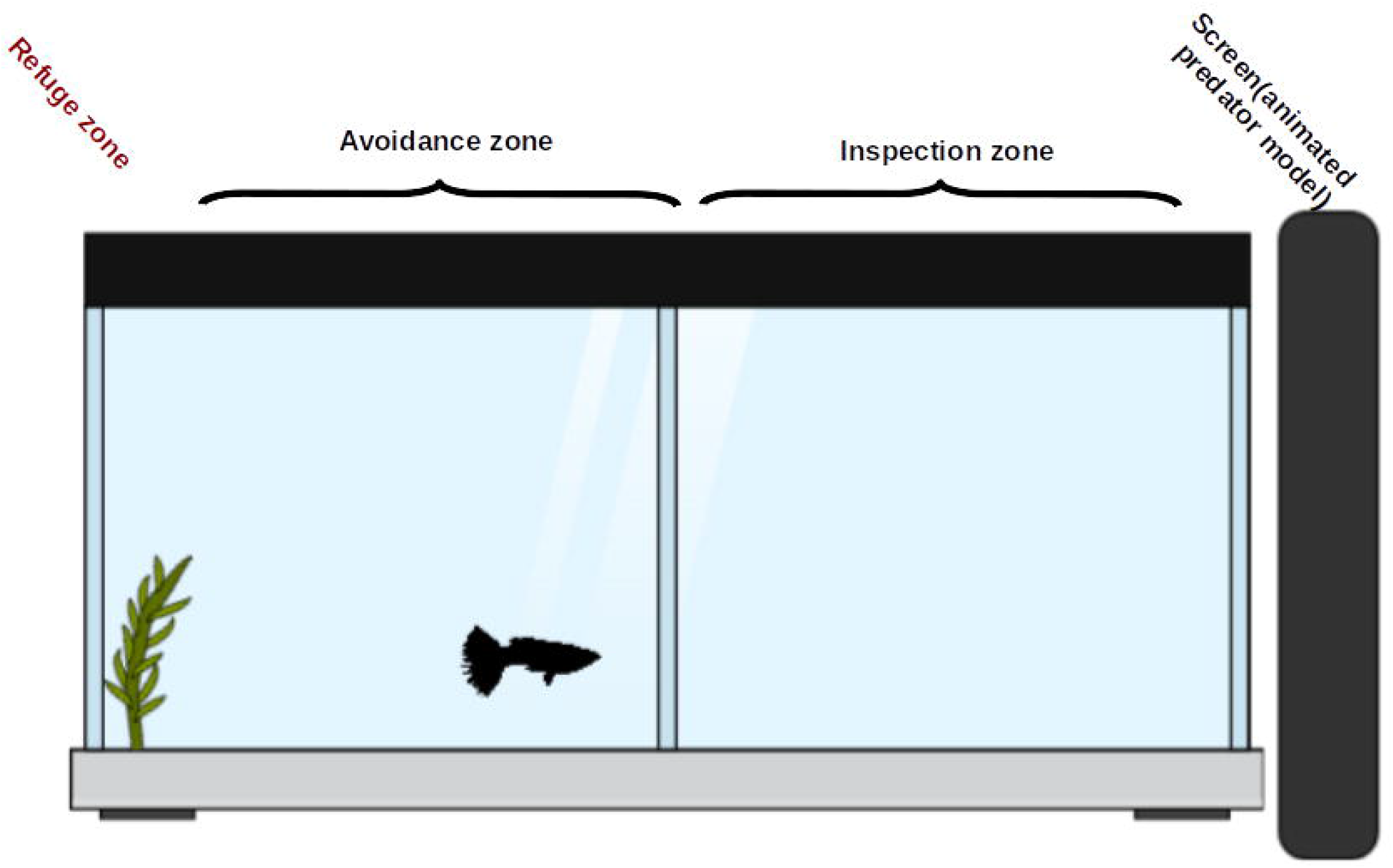

Conditional approach was induced by playback of an animated predator (blue acará cichlid *Aequidens pulcher*), produced and presented using LibreOffice Impress (Version: 5.3.7.2) on a CPU running Linux 4.15 (Xubuntu 16.04).The animation was presented at the computer screen for the last 6 min of each session, with the exception of the “Predator absent” condition of Experiment 1. The animals are expected to respond to acará cichlids with antipredator behavior, which has been observed in other populations (e.g. ref. [34]).

### 2.3. Open science practices

The experiments were not formally pre-registered. Datasets and scripts are available from https://github.com/lanec-unifesspa/TFT (doi: 10.5281/zenodo.1290044).

### 2.4. Experiment 1

#### 2.4.1. Experimental design, endpoints, and statistical analysis

In the first experiment, 16 animals were used, divided equally and randomly between two groups: “Predator absent” and “Predator present”. For both groups, the mirror was positioned in parallel to the side of the tank. In the first group, animals were introduced to the tank and allowed to explore it for 10 min; in the 5^th^ minute, the video was not turned on. The second group was subjected to the same manipulations, except that the video was turned on in the 5^th^ minute. The order of subjects was randomized using http://randomization.com/. Video recordings of the behaviour were made and later analyzed with manual event recording, with the help of the software X-Plo-Rat 2005 (https://github.com/lanec-unifesspa/x-plo-rat [35]) by two trained observers. Observers were not blind to group allocation, as it was not possible to exclude the portion of the video that included the monitor. Times spent in the avoidance, refuge, or inspection zone were computed as percent change, as follows:

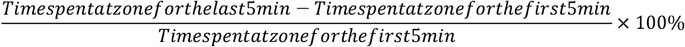

These values were compared using independent 2-group Mann-Whitney’s U tests, with *p*-values < 0.05 considered statistically significant. Effect sizes for these variables were reported as Cohen’s *d*. Raw values for the acclimation period were also analyzed, to understand whether there were biases or baseline differences across groups that could be responsible for possible effects.

Data on freezing was analyzed as raw values, in s, of the time spent freezing. These data were analyzed using a repeated measures ANOVA, with time block as within-subjects effect, and group as between-subjects effect. Effect sizes for freezing were reported as ω^2^.

### 2.5. Experiment 2

#### 2.5.1. Drug treatments

Fluoxetine hydrochloride (CAS #54910-89-3) was bought from Libbs. Metergoline (CAS #17692-51-2) was bought from Virbac Brasil. All treatments were acute – i.e. a single injection was made, 20 min. before experiments. The fluoxetine dose was based on previous work with zebrafish [36], which showed an antipanic-like effect of this dose in an acute treatment. The metergoline dose was based on previous work with rats [37], which showed an effect of this dose on the forced swimming test.

Injection procedure was adapted from a protocol existing for intraperitoneal administration in zebrafish [38]. Briefly, animals were anesthetized in cold water (12 °C – 17 °C), and transferred to a “surgical bed” made of a washing sponge with a through that allowed positioning the animal with the ventral side exposed, while simultaneously perfusing the gills. The sponge was soaked with ice-cold water to maintain gill perfusion, and the setup was kept on ice. Drugs were injected at a volume of 2 μL/animal, using a Hamilton microsyringe (701N, needle gauge 26s), and animals were immediately transferred to aerated water at room temperature to recover.

#### 2.5.2. Experimental design and statistical analysis

In the second experiment, 30 animals were used, divided equally and randomly between three groups: a control group (CTRL), injected with vehicle (Cortland’s salt solution; https://wiki.zfin.org/display/prot/Cortland%27s+salt+solution); a fluoxetine-treated group (FLX), injected with 2.5 mg/kg fluoxetine; and a metergoline-treated group (MET), injected with 1 mg/kg metergoline. The dose was based on the average weight of the animals. Experimenters were blinded to treatment by using coded vials for drugs. 20 min after injection, animals were individually transferred to the test tank, and subjected to the same manipulations as the EXP group from Experiment 1. The order of subjects was randomized using http://randomization.com/. Data were recorded and registered as described in Experiment 1, with spatiotemporal endpoints (time spent in avoidance, refuge, and inspection zones) indexed as percent change, and data for freezing analyzed raw. Spatiotemporal data was analyzed using Kruskal-Wallis’ rank sum test followed by Dunn’s post-hoc multiple comparison tests adjusted with the Benjamini-Hochberg method; freezing data was analyzed using a repeated measures ANOVA, with time block as within-subjects effect, and treatment as between-subjects effect. Effect sizes for freezing were reported as ω^2^.

## 3. Results

### 3.1. Experiment 1

No differences were found in baseline (pre-animation) values across groups on time in the refuge (W = 23, p = 0.3706; data not shown), avoidance zone (W = 29, p = 0.7984; data not shown), or inspection zone (W = 40, p = 0.4418; data not shown). The animated predator increased time in the inspection zone when the mirror was parallel to the tank (Figure 2A; W = 13, p = 0.0498, *d* = −1.25). The animated predator did not change time spent in the avoidance zone (Figure 2B; W = 24, p = 0.4418, *d* = −0.49). Refuge use was altered by the animated predator (Figure 2C; W = 59, p = 0.003, *d* = 0.82). Finally, the animated predator increased freezing in the second time block (i.e. after the stimulus was turned on), but not in the first time block (Figure 2D; F_[1,28]_ = 6.46, p = 0.0169, ω^2^ = 0.138 for the main effect of group).

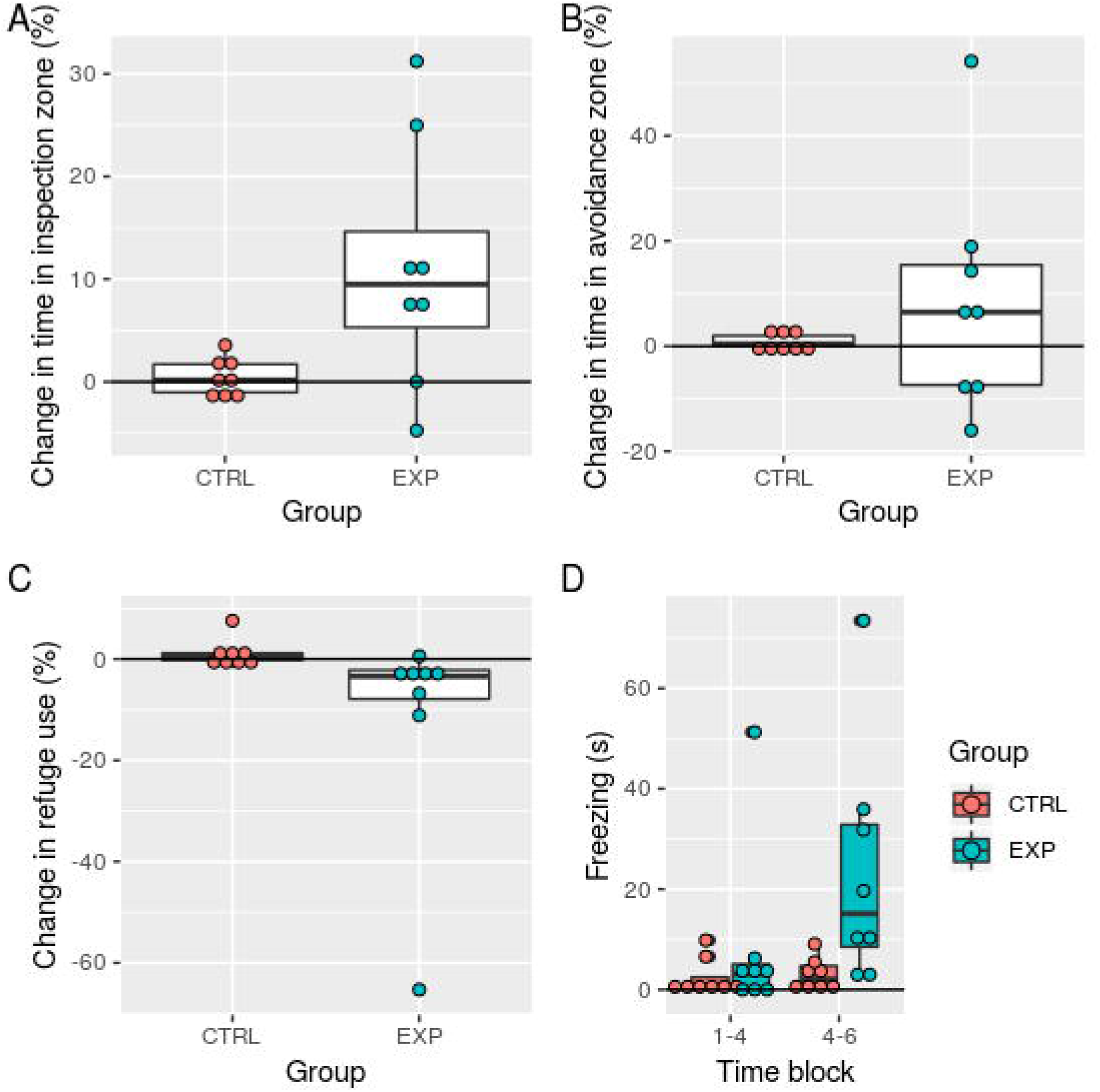

### 3.2. Experiment 2

Acute fluoxetine increased time spent in the inspection zone (Z = −2.16, p = 0.046 vs. control), while metergoline decreased time in the inspection zone (Z = 2.01, p = 0.045 vs. control)(Figure 3A; *χ*^2^_[df = 2]_ = 17.36, p = 0.00017, ω^2^ = 0.476). Only fluoxetine produced an effect on time spent in the avoidance zone (Figure 3B; ω^2^_[df = 2]_ = 16.059, p = 0.00033, ω^2^ = – 0.555), with fluoxetine decreasing time spent in the avoidance zone (Z = 2.616, p = 0.013 vs. control), and metergoline having no effect (Z = −1.32, p = 0.186 vs. control). Neither fluoxetine nor metergoline altered refuge use (Figure 3C; *χ*^2^_[df = 2]_ = 1.319, p = 0.517, ω^2^ = – 0.00331). Freezing was increased by fluoxetine (p = 4.0·10^-7^ vs. control), while metergoline decreased it (p < 4.0·10^-7^ vs. control)(Figure 3D; F_[2,27]_ = 26, p = 5.2·10^-7^, ω^2^ = 0.255 for the interaction term).

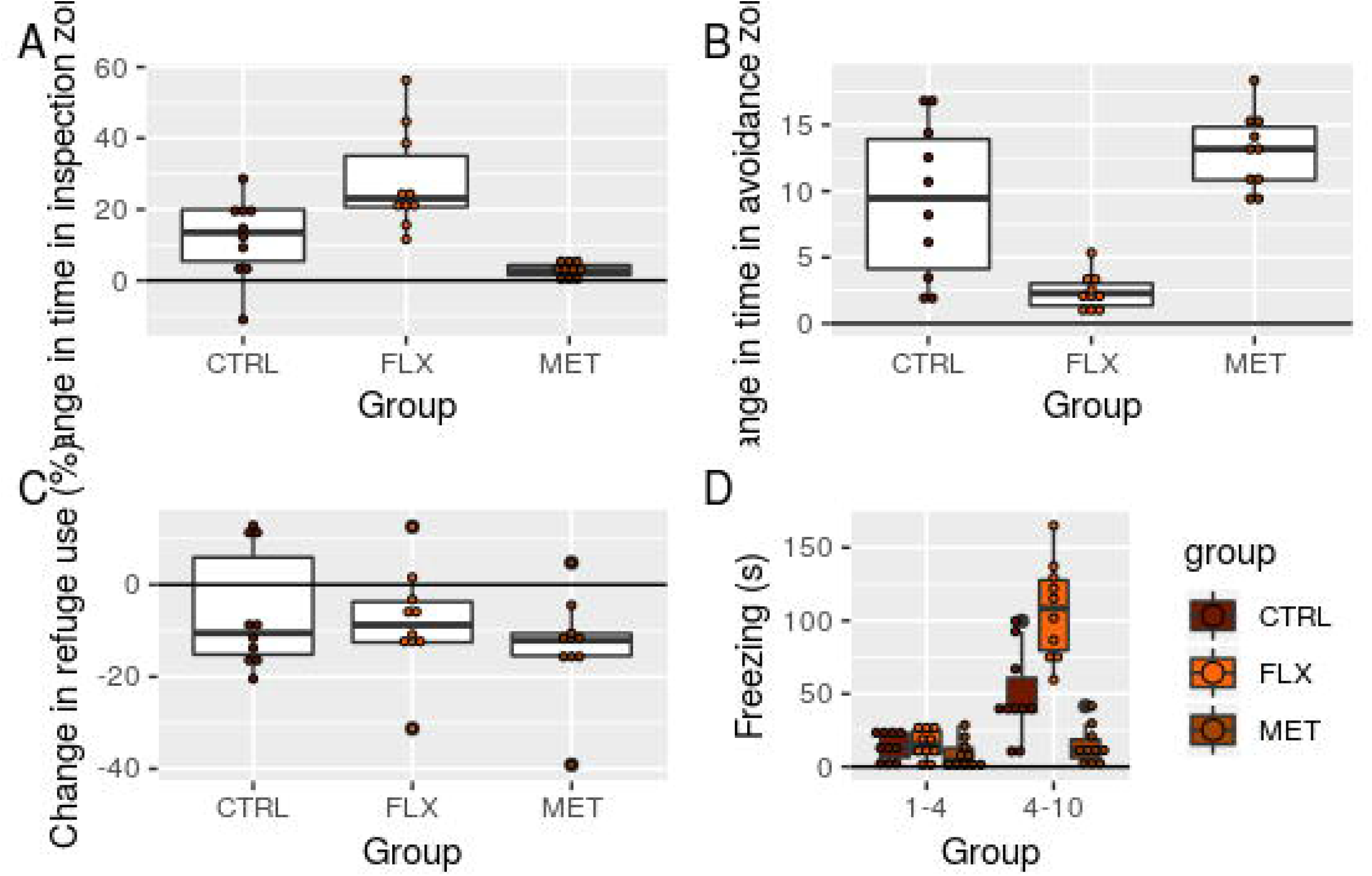

## 4. Discussion

In the present work, we replicated the observation made by Dugatkin [5,6]on conditional approach by guppies during predator inspection, and extended his findings by showing that guppies also freeze more during predator inspection. We also show that phasic serotonin promotes conditional approach/cooperation but increases freezing/fear, and limited evidence was found for a tonic participation of serotonin in conditional approach.

We predicted that the animated predator image would promote predator inspection, but would also increase fear, reflected by increased freezing, as well as time spent in the avoidance and refuge zones. While freezing and refuge use were indeed increased, animals did not appear to avoid the predator. We suggest that freezing, in this situation, is an optimal strategy to avoid being detected by the predator while at the same time benefiting from gathering information, maintaining a high level of alertness. The relationship between fear, stress, and social behaviour has seldomly been addressed in the literature [27,39]. There is some evidence from cleaning gobies *Elacatinus* spp. that cooperating in cleaning mutualisms with piscivorous clients (potential predators) is stressful, as cortisol levels are increased during these interactions [40]. Interestingly, field experiments in cleaner wrasse showed that treament with cortisol increases “cheating”, in which cleaners provide small clients with more tactile stimulation with their pectoral and pelvic fins – a behaviour that attracts larger clients that are then bitten to obtain mucus –, while blocking glucocorticoid receptors led cleaners to increase tactile stimulation to larger clients [41].

An important concern of our experiments is that, differently from Dugatkin [5], the animals used in Experiments 1 and 2 are not wild-caught, but laboratory-reared, and the stimulus predator was the blue acará cichlid. As a result, animals could have been displaying behavior that is not specifically anti-predator in nature, but instead represent a “curiosity” approach to any novel fish stimulus. Indeed, in wild populations from Trinidad, pike cichlids (*Crenicichla lepidota*) induce a “surfacing” behavior that is not observed when animals are exposed to acará cichlids [34]; however, even in wild populations, acará and pike cichlids do not differ in their ability to induce an inspection response or in their ability to induce fear-like behavior (e.g. inhibit foraging and increase shoaling) [34]. While wild-caught female guppies from high-predation areas appear to respond less to blue acará cichlids than to pike cichlids *C. frenata*, this difference disappears at the first and second generations derived from these wild populations [42]. In the light of these findings, and considering the effect that our blue acará cichlid animation had on freezing, our results suggest that this domesticated population is displaying normal antipredator behavior, and not simply a novelty response. However, a neophobic response cannot be fully discarded as driving part of the behavior reported here.

An alternative explanation to the results from Experiment 1 is that predator inspection does not represent conditional approach, but instead males undertake inspection as a demonstration to females of their superiority in relation to non-inspectors. Indeed, it has been demonstrated that female guppies prefer bolder males, which show higher levels of inspection [43,44], and bolder individuals tend to produce more social interactions [45]; assortative interactions and sexual selection are expected to play a role in the establishment of this strategy [8]. However, this is not necessarily contradictory with the conditional approach strategy since, differently from “pure” tit-for-tat conditional approach does not require some form of kin or group selection to be at work. Conditional approach can be better understood in a “social competence” approach [12,46], in which situation-specific cues (such as presence or absence of females, or varying risk levels), together with various other factors, allow individuals to assess the situation in order to make behavioral decisions. While risk taking/boldness certainly takes part on this [4], it is not the only variable responsible for the development of predator inspection in our paradigm.

In Experiment 2, we hypothesized tonic and phasic roles for serotonin in promoting cooperation and increasing fear. We found that acute fluoxetine increased inspection and fear, while acute metergoline decreased both. Guppies under treatment with fluoxetine spent more time in the inspection zone (i.e. nearer to the image of a predator) and less in the avoidance zone. However, fluoxetine also increased freezing, suggesting that its effects were not fully explained by simply increasing “boldness” (or, conversely, decreasing fear and/or anxiety). In the cleaner wrasse, fluoxetine increased the probability of cleaning clients without altering cleaning quality [31], suggesting that phasic serotonin increases the motivation to cooperate in fish [47]. Differently from the cleaner wrasse, however, conditional approach during predator inspection entails exposure to threat, while most of the clientele of cleaners is harmless. Acute fluoxetine increased freezing, while metergoline decreased it, suggesting that serotonin controls antipredator behaviour in guppies. Similar observations have been made in zebrafish, in which serotonin appears to have a “dual role” in mediating responses to potential vs. real threat [25].

While phasic serotonin appears to motivate cooperation in guppies, we found evidence of a serotonergic tone promoting conditional approach. This is consistent with other findings from the literature: tryptophan depletion reduces cooperation in the Prisoner’s Dilemma in human participants [30], and 5-HT_1A_ receptor antagonism or eliminating 5-HT tonus decreases cleaning in cleaner wrasses [31]. In this last case, reducing 5-HT activity also decreased cheating frequencies due to an overall reduction of the proportion of clients inspected and of the average duration of interactions [31]. These effects may occur via mediation of risk perception, which would enhance the cleaner’s appraisal of the threat represented by predatory clients [27,39,47,48].

## 5. Conclusion

Overall, the results reported present a scenario in which serotonin acts to promote cooperation in a laboratory setting in fish. These results contribute to the description of the neurochemical bases of pro-social behaviour, and could have translational applications. Moreover, they also shed light on the relationship between fear and sociality, an area seldomly explored in the social behaviour literature [27,39]. Whether this is also modulated by mediators of fear and stress, such as cortisol, is still unknown.

## Acknowledgments

AFNP is the recipient of a CNPq/PIBITI scholarship. CM is the recipient of a CNPq/Universal grant (#400726/2016-5). MCS is the recipient of two FCT grants (SFRH/BPD/109433/2015; PTDC/MAR/105276/2008).

